# Brain Serotonergic Fibers Suggest Anomalous Diffusion-Based Dropout in Artificial Neural Networks

**DOI:** 10.1101/2022.05.22.492968

**Authors:** Christian Lee, Zheng Zhang, Skirmantas Janušonis

## Abstract

Random dropout has become a standard regularization technique in artificial neural networks (ANNs), but it is currently unknown whether an analogous mechanism exists in biological neural networks (BioNNs). If it does, its structure is likely to be optimized by hundreds of millions of years of evolution, which may suggest novel dropout strategies in large-scale ANNs. We propose that the brain serotonergic fibers meet some of the expected criteria because of their ubiquitous presence, stochastic structure, and ability to grow throughout the individual’s lifespan. Since the trajectories of serotonergic fibers can be modeled as paths of anomalous diffusion processes, in this proof-of-concept study we investigated a dropout algorithm based on the superdiffusive fractional Brownian motion (FBM). This research contributes to biologically-inspired regularization in ANNs.

## 1 INTRODUCTION

Random dropout is a simple but powerful technique employed in the training of artificial neural networks (ANNs). Its main goal is to improve network regularization and minimize overfitting (Srivastava et al., 2014; Goodfellow et al., 2016). In the standard implementation, the output of a randomly selected set of hidden units is set to zero, and this functional elimination is repeated in each training iteration. The eliminated units neither participate in the current forward pass nor contribute to the backpropagation calculations. As a consequence, the network has a slightly different architecture with each input and cannot heavily rely on any individual neurons (Hinton et al., 2012; Krizhevsky et al., 2012). Conceptually, this technique can be thought of as an efficient approximation of bagging, in which a set of different models is trained on a shared dataset (Goodfellow et al., 2016). In ANNs, the dropout rate is typically 10-50%; the higher rates (40-50%) are common in convolutional neural networks (CNNs) (Geron, 2019).

ANNs are fundamentally different from biological neural networks (BioNNs), just as the artificial and biological neurons have little in common. In fact, direct mimicking of BioNNs can be counterproductive, as exemplified by the relatively recent transition from the “more natural” sigmoid activation function and to the less biological but more efficient rectified linear activation function (ReLU) (Krizhevsky et al., 2012). On the other hand, BioNNs are superior to ANNs in their highly sophisticated level of abstraction, which allows them to robustly learn from very small training sets (sometimes a single instance).

The development of ANNs is primarily motivated by practical applications, not by neuroscience. However, each of the two fields has stimulated novel insights in the other. For example, the fundamental architecture of modern CNNs has deep roots in the visual neuroscience of the early 1960s and, in particular, in the notion of a hierarchical system of “receptive fields” (Hubel and Wiesel, 1962; LeMasurier and Van Wart, 2012). On the other hand, recent advances in CNNs have convincingly demonstrated that complex neuroanatomical circuits with many specialized regions (Sporns, 2010) or large-scale oscillations (He et al., 2010) are not necessary for reliable detection and segmentation of objects in complex visual scenes or for human speech recognition (Alzubaidi et al., 2021). Presently, a major effort is underway to make ANNs more “intelligent” (by getting closer to the brain’s ability to operate at a high level of abstraction), which has led to the development of a benchmark dataset, named the Abstraction and Reasoning Corpus (ARC) (Chollet, 2019).

Dropout is peculiar in that it is now a standard and well-validated method in ANN training but it has no obvious counterpart in the biological brain. Logic suggests that BioNNs can also overfit, at the expense of deeper abstractions (as perhaps manifested in savant memory (Bor et al., 2007; Song et al., 2019)). If a dropout-like mechanism actually exists in neural tissue, it can be expected to be (*i*) ubiquitously present and have a structure that is both (*ii*) strongly stochastic and (*iii*) unstable at the level of individual neurons, including the adult brain. These requirements may be met by the serotonergic fibers, a unique class of axons described in virtually all studied nervous systems (vertebrate and invertebrate).

In vertebrates, the serotonergic fibers are axons of neurons located in the brainstem raphe complex (Jacobs and Azmitia, 1992; Okaty et al., 2019). These fibers travel in extremely long, meandering trajectories and form dense meshworks in virtually all brain regions (Steinbusch, 1981; Lidov and Molliver, 1982; Foote and Morrison, 1984; Lavoie and Parent, 1991; Vertes, 1991; Voigt and de Lima, 1991; Morin and Meyer-Bernstein, 1999; Vertes et al., 1999; Linley et al., 2013; Migliarini et al., 2013; Donovan et al., 2019). Early estimates have suggested that each cortical neuron in the rat brain is contacted by around 200 serotonergic varicosities (dilated fiber segments) (Jacobs and Azmitia, 1992). The electrophysiological characterization of serotonergic neurons remains grossly incomplete, given their diversity (Okaty et al., 2019). Early studies have reported neurons that fire at remarkably stable rates (Jacobs and Azmitia, 1992), suggesting low information transmission. More recent research has shown that some serotonergic neurons respond to conditions that require learning in uncertainty (Matias et al., 2017), and that serotonin (5-hydroxytryptamine) is fundamentally associated with neural plasticity (Lesch and Waider, 2012). The renewed interest in therapeutic applications of serotonin-associated psychedelics is motivated by the recent findings that these psychedelics can be surprisingly efficient in rapidly boosting cognitive flexibility – thus opening up new opportunities in the treatment of brain disorders associated with cognitive persistence (Vollenweider and Preller, 2020; Daws et al., 2022). Conceptually, serotonin may support “unfreezing” of synapses that may have become “locked in” or “overfitted.”

In addition, recent research has shown that the trajectories of serotonergic fibers are strongly stochastic. Therefore, the number of fiber contacts received by an individual neuron in any brain region is a random event. The mathematical models of serotonergic trajectories are an active area of research (Janušonis and Detering, 2019; Janušonis et al., 2020). Some features of these fibers are captured by the superdiffusive fractional Brownian motion (FBM), an anomalous diffusion process that generalizes normal Brownian motion. FBM is self-similar stochastic process parametrized by the Hurst index (*H*, where 0 < *H* < 1) (Mandelbrot and Van Ness, 1968). In the superdiffusive regime (*H* > 0.5), increments are positively correlated over long times, which endows these paths with *H*-dependent “persistence” (Janušonis et al., 2020; Vojta et al., 2020).

Importantly, the paths of individual serotonergic fibers may continuously change, also in the adult brain. Experimental research has demonstrated that serotonergic fibers are nearly unique in their ability to robustly regenerate in the adult mammalian brain after an injury, and that regenerating fibers do not follow their previous paths (Jin et al., 2016; Cooke et al., 2022). Long-term live imaging of serotonergic fibers in intact brains currently poses major technical challenges. However, circumstantial evidence suggests that serotonergic fibers may undergo routine regeneration in the healthy brain. They are extremely long, thin, and not fasciculated, which may result in frequent interruptions because of local tension forces and biological processes, such as microglial activity (Janušonis et al., 2019). This dynamic would continuously generate new fiber paths beyond the interruption points.

In summary, serotonergic fibers have a number of features that are conducive for a dropout-like mechanism in biological neural tissue. In one scenario, serotonergic fiber contacts may interfere with the normal activity of individual neurons, effectively removing them from the network. Alternatively, these contacts can stabilize the output of individual neurons. *In vitro*, the growth rate of serotonergic axons can be remarkably fast, with long extensions over the course of hours (Azmitia and Whitaker-Azmitia, 1987). To our knowledge, no reliable *in vivo* growth rate estimates are currently available in the healthy adult brain.

In this study, we examined an FBM-based dropout algorithm in simple ANNs. We show that the performance of this dropout is comparable to that of the standard dropout. At the same time, it is considerably more biologically realistic and may stimulate further investigations of its efficiency in complex, large-scale network architectures.

## 2 METHODS

An FBM-based dropout method (further referred to as the “FBM-dropout” was tested in a fully connected network consisting of an input layer (with the ReLU activation function), a hidden layer (with the ReLU activation function), and an output layer. The hidden layer had *N×N* neurons and was endowed with the Euclidean geometry, in addition to the standard topological structure. Its neurons were arranged in a square grid extending from 0 to 1 units in both dimensions. In the grid, each neuron was represented by a square with the side of 1/(2*N*) units, and adjacent neurons were spaced by the same distance (1/(2*N*) units) in both dimensions.

Two-dimensional FBM paths were generated using the Python *stochastic* (0.6.0) package. The two coordinates were modeled as independent one-dimensional paths (with the same *H*, the drift μ = 0, and the volatility σ = 1). The value of *H* was set at 0.9 based on previous experimental research (Janušonis et al., 2020). For each training epoch, a number (*n*) of FBM paths were generated in the time interval [0, *T*], and their numerical values at each Δ*t* time-units were stored in an array (as the entire trajectory of each fiber). The variable *n* was used to control the average dropout rate (it can also be controlled by adjusting the “length” of the fibers or the geometry of the neurons, but these approaches are not equivalent). Based on an empirical optimization, in the presented analyses we set *T* = *i*_max_/10 and Δ*t* = 1/500, where *i*_max_ is the maximal number of training iterations (the actual number of iterations may be lower). We note that *T* and Δ*t* refer to the FBM process itself and are not directly related to the network training “time”; the two can be flexibly coupled.

In each iteration (*i* = 0, 1, 2,…), each fiber was modeled as a sliding subarray representing the time interval [(*i*×*s*)×Δ*t*, (*i*×*s*+*L*)×Δ*t*], where *L* and *s* are positive integers representing the fiber “length” (in the number of points) and the shift (in the number of points), respectively. Note that if *s* = *L*, the fiber advances fast and in each iteration starts where it ended in the previous iteration. If *s* < *L*, the fiber “crawls” more slowly, advancing fewer steps and retaining some of its previously occupied positions. In the presented analyses, we set *L* = *s* = 50.

Extremely long FBM paths are computationally expensive and often require supercomputing resources (Janušonis et al., 2020; Vojta et al., 2020), due to their long-range dependence on all previous steps (if *H* ≠ 0.5). To simplify computations, we assumed that at the end of each training epoch the fiber “branches,” initiating a new FBM path at a random point of the last segment, also accompanied by the instant “degeneration” of the previous path. In order to avoid boundary effects (which would require modeling reflected FBM paths (Janušonis et al., 2020; Vojta et al., 2020) but would not be biologically meaningful here), we implemented the periodic boundary conditions (i.e., the fiber never leaves the layer and reemerges on the opposite side when it crosses a border).

The dropout was modeled as follows: if any of the fibers crossed the boundary of a neuron, the output of this neuron was set to zero. There was no interaction among the fibers (*e*.*g*., the same neuron could be contacted by more than one fiber). The described model is a simplification of biological reality, where serotonergic fibers are always attached to a cell body and may instead continuously regenerate with new paths from random interruption points (*i*.*e*., they do not actually “crawl” as detached segments). However, the overall dynamic of the model does approximate these biological processes, including axon branching.

All ANN training and testing scripts were written in Python 3 with the PyTorch package (1.11.0).

## 3 RESULTS

The standard dropout and the FBM-dropout are fundamentally different in that in the former case neurons are turned off independently of other neurons, but in the latter case neurons in close proximity are more likely to be turned off at the same time (because they are more likely to fall under the same “slithering” fiber). This introduces spatial correlations, which can be tightened or relaxed by controlling the number and “length” of the fibers, their FBM parameter (*H*), their “speed” (*s*), and the geometry of the neurons (*e*.*g*., they can be sparsely or densely packed). Despite this more structured dropout, all neurons can get visited at some time during the training (Fig. 1). In addition, the overall trajectories of the fibers resemble the trajectories of actual serotonergic fibers in brain tissue (Fig. 1A and F). The iterative dynamics of multiple fibers is shown in Figure 2.

**Figure 1.**
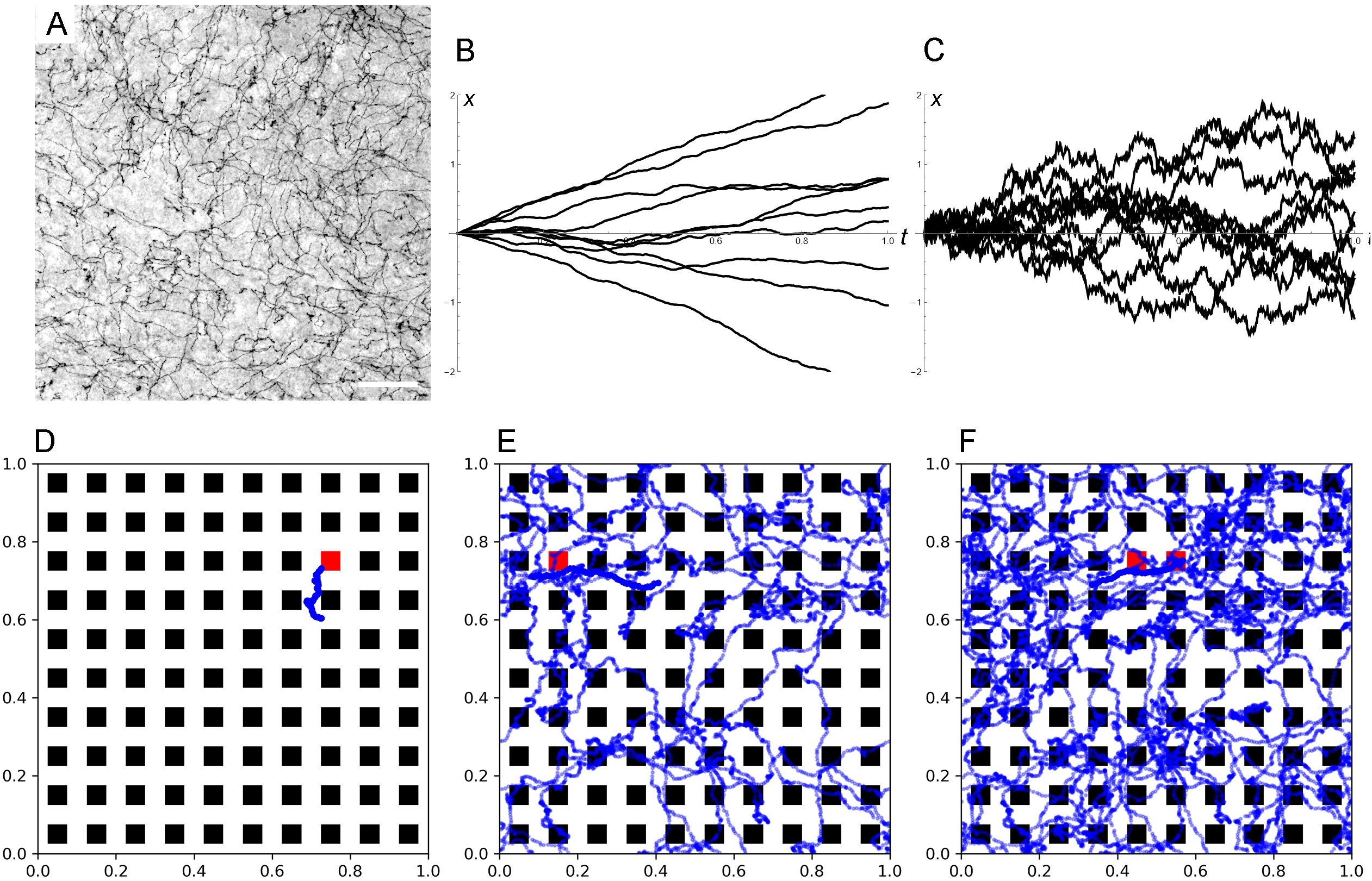
(A) Serotonergic fibers visualized with immunohistochemistry for the serotonin transporter in the mouse primary somatosensory cortex. This bright-field image represents three focal levels (in a 40 μm-thick coronal section) that have been digitally merged. Scale bar = 50 μm. The immunohistochemistry was performed as described in (Janušonis et al., 2020). (B) Ten sample paths of one-dimensional FBM with *H* = 0.9 (μ = 0, σ = 1, *T* = 1). (C) Ten sample paths of one-dimensional FBM with *H* = 0.5 (μ = 0, σ = 1, *T* = 1), corresponding to normal Brownian motion. Normal Brownian motion is not appropriate for the modeling of serotonergic fibers, but it plays a central role in the theory stochastic processes (*e*.*g*., it is used to model simple diffusion and financial markets). It is shown here for comparison. (D-F) The history of a *single* simulated fiber traveling in a hidden layer with 100 neurons (small squares), shown at 1 (D), 150 (E), and 300 (F) training iterations. The current position of the fiber is represented by a thick, solid segment. In actual training, more than one fiber is used. The neurons that are dropped out at each step are colored red. *H* = 0.9, *T* = 30, Δ*t* = 1/500, *L* = 50, *s* = 50.

**Figure 2.**
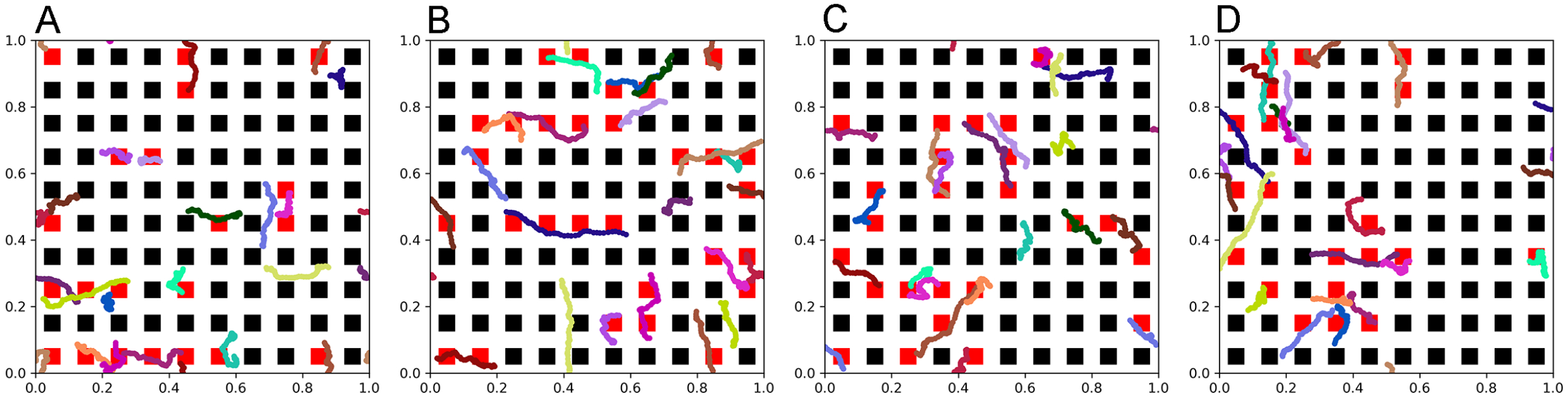
The dynamics of a set of fibers (randomly color-coded) at 75 (A), 150 (B), 225 (C), and 300 (D) iterations. The neurons that are dropped out at each step are colored red. All fibers are the same “length” in the sense that they represent equal time intervals in the FBM path. (FBM paths are fractal and do not have a defined length in the usual sense.) *H* = 0.9, *T* = 30, Δ*t* = 1/500, *L* = 50, *s* = 50.

We examined the performance of the network (with regard to overfitting) in a simple training example. A random set of 50 points in the range of [-1, 1] was generated, and a linearly-dependent second set of points (*y*) was produced, with an additive noise term (*y* = *x*+ 3ε, where ε has a normal distribution with the mean of 0 and the standard deviation of 1). A network with 100 neurons in the hidden layer was trained on this set with no dropout, the standard dropout, and the FBM-dropout. The FBM-dropout strongly outperformed the no-dropout condition (where pronounced overfitting was observed, as expected) and was indistinguishable from the standard dropout (Fig. 3).

**Figure 3.**
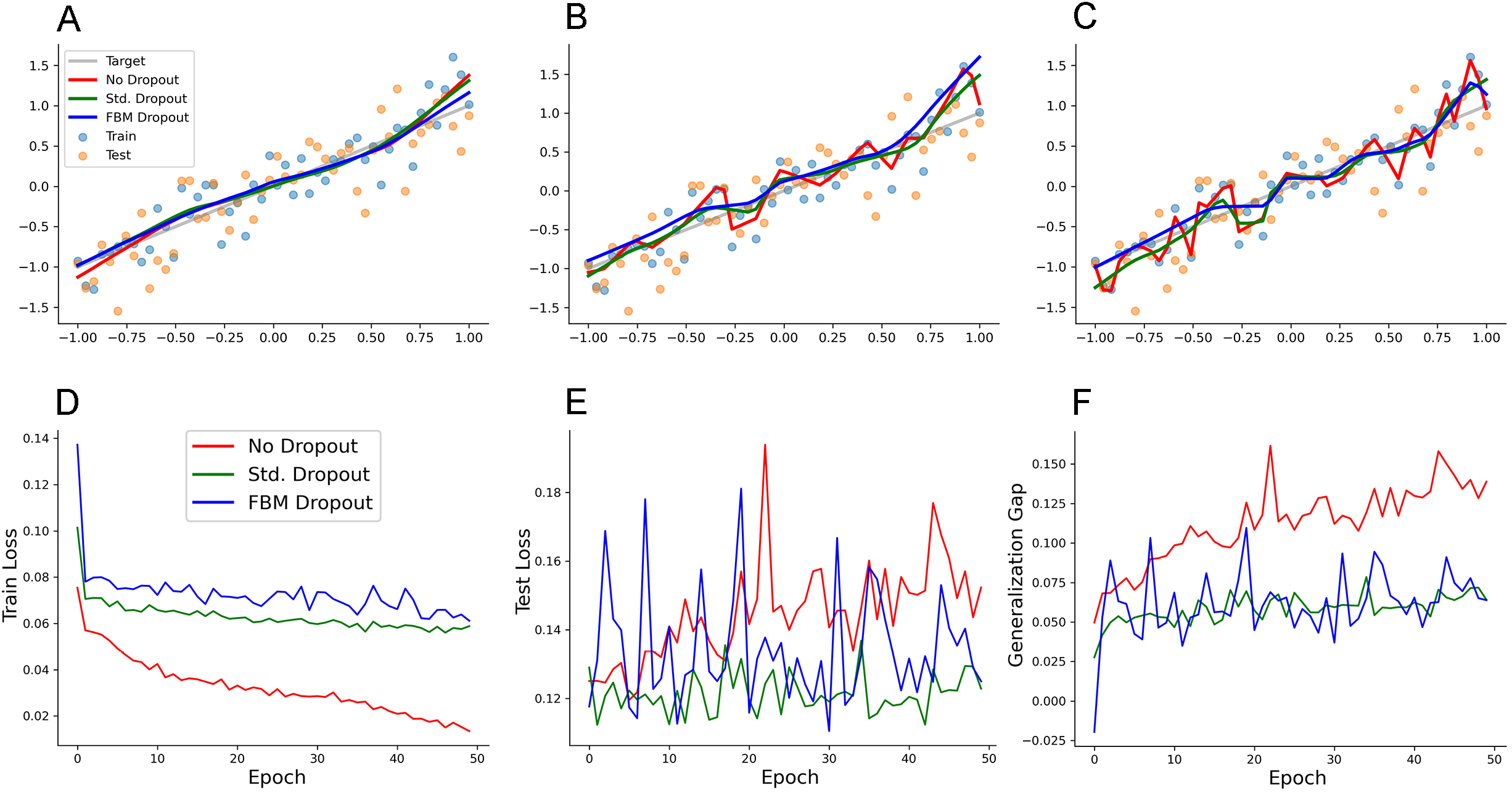
A regression-type model trained with no dropout (red in all panels), with the standard dropout (green in all panels), and with the FBM-dropout (blue in all panels). (A-C) The fitted curves at 1 (A), 20 (B), and 50 (C) epochs. (D) The training loss over the training epochs. (E) The testing loss after each training epoch. (F) The generalization gap (the difference between the testing loss and the training loss) after each training epoch. The network consisted of one input neuron, 100 neurons in the hidden layer, and one output neuron. The number of epochs was 50, with 50 mini-batches with 1 sample each. The Adam optimizer with the learning rate of 0.01 was used. In all conditions, the dropout probability was adjusted to be around 0.2. *N* = 10, *n* = 12, *H* = 0.9, *T* = 5, Δ*t* = 1/500, *L* = 50, *s* = 50. The same dropout parameters were used in the input and hidden layers (both with independent sets of fibers, which stayed within their respective layers).

We next examined the performance of the network on a reduced set of the MNIST hand-written digits (http://yann.lecun.com/exdb/mnist/). In the reduced set, 1000 training samples and 1000 testing samples were randomly selected from the original set. A network with 1024 neurons in the hidden layer was trained on this set with no dropout, the standard dropout, and the FBM-dropout. The FBM-dropout again performed well compared to the standard dropout (Fig. 4).

**Figure 4.**
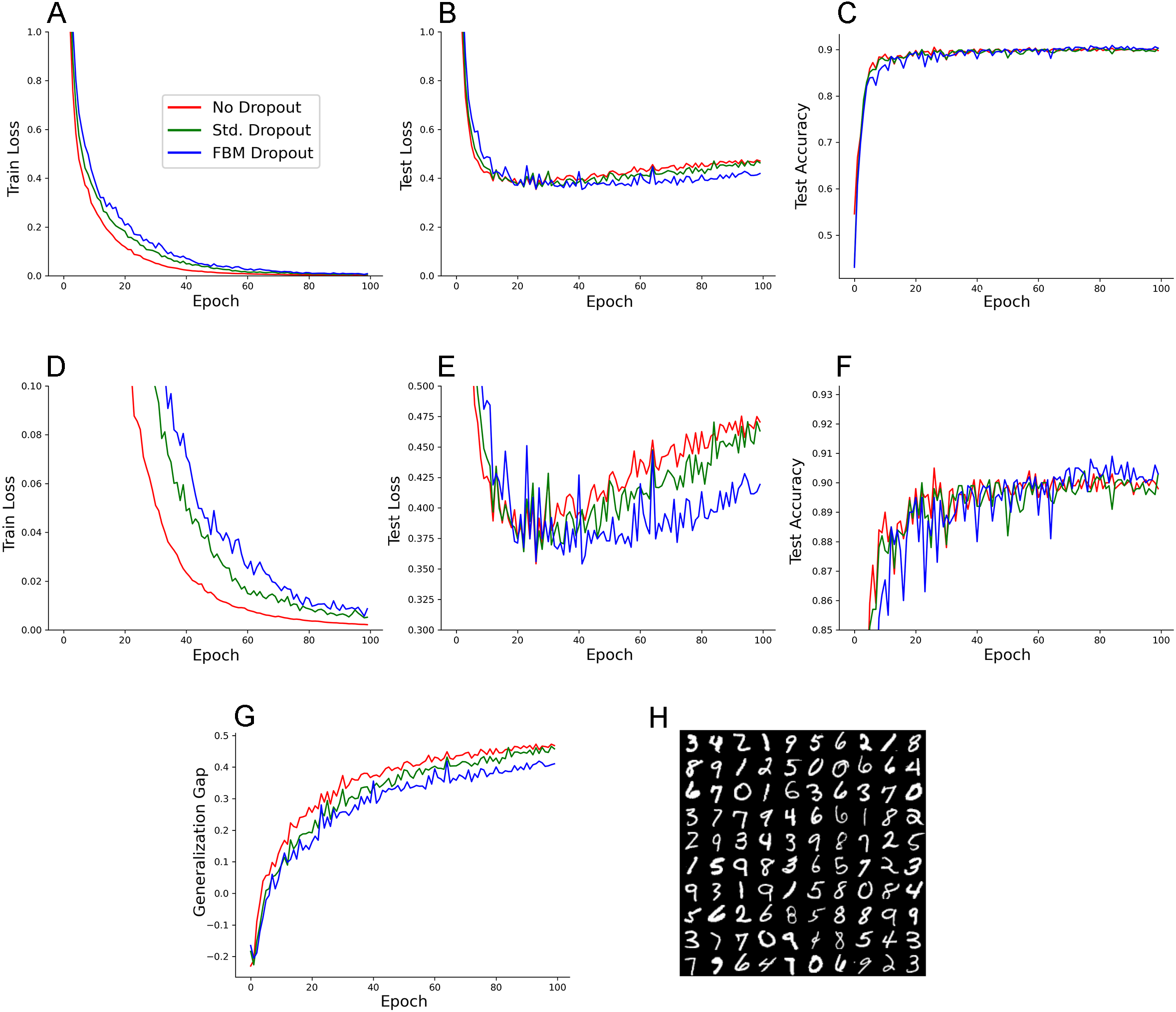
An MNIST model trained with no dropout (red in all panels), with the standard dropout (green in all panels), and with the FBM-based dropout (blue in all panels). (A) The training loss over the training epochs. (B) The testing loss after each training epoch. (C) The testing accuracy after each training epoch. (D-F) A zoomed-in view of the panels above. (G) The generalization gap (the difference between the testing loss and the training loss) after each training epoch. The network consisted of 784 input neurons (28 × 28 grayscale images of digits), 1024 neurons in the hidden layer, and 10 output neurons. The number of epochs was 100, with 16 mini-batches with 64 samples each. The Adam optimizer with the learning rate of 0.0001 was used. In all conditions, the dropout probability was adjusted to be around 0.2. *N* = 32, *n* = 60, *H* = 0.9, *T* = 2, Δ*t* = 1/500, *L* = 50, *s* = 50. The same dropout parameters were used in the input and hidden layers (both with independent sets of fibers, which stayed within their respective layers). (H) Examples of the images used in the training and testing.

## 4 DISCUSSION

Dropout was introduced around ten years ago (Hinton et al., 2012; Labach et al., 2019) and since has become a standard technique in the machine learning field. Despite the computational simplicity and effectiveness of random dropout in some ANNs, it has serious limitations in important network architectures. These networks include the powerful CNNs, where random dropout has little effect due to the highly correlated pixels in feature maps (Labach et al., 2019). As ANNs become larger and more complex in their architecture, dropout algorithms are likely to evolve in several directions.

Here, we present an approach that is strongly motivated by neurobiology and is built on recent analyses of serotonergic fibers, led by one of the co-authors (Janušonis and Detering, 2019; Janušonis et al., 2020). We demonstrate the feasibility of this approach in simple, proof-of-concept networks, where it performs at least as well as random dropout. However, it has a rich statistical structure which may serve as a toolbox for future improvements of dropout techniques.

Conceptually, the method is simple: the relevant neuron layers are placed in a Euclidean space and enriched with fiber-like entities that move through this space. When a fiber comes into contact with a neuron, the neuron becomes (temporarily) inactive. Computationally, a number of parameters can be easily adjusted, resulting in different dropout statistics. These parameters include the geometry of the layer (the size and shape of the neurons, as well as their spacing which can be deterministic or stochastic) and the fibers themselves, which can differ in their numbers, *H* values, “length,” and “speed.” For example, many short, fast moving fibers with *H* = 0.5 will approximate random dropout, but one long, slow moving fiber with *H* > 0.5 will result in strongly correlated dropout events.

An intriguing extension of this method would be adding a third dimension and allowing fibers to move across network layers, as serotonergic fibers do in brain tissue. Tracing studies have shown that a single serotonergic fiber can traverse multiple brain regions, separated by vast anatomical distances (Gagnon and Parent, 2014). This extension is not trivial conceptually, given the topological nature of ANNs, but it may lead to interesting findings. Computationally, it would produce correlated dropout events at different processing levels in the network hierarchy, which might be beneficial in CNNs. It may also find applications in artificial spiking neural networks (SNNs) which already encode spatial and temporal information (Pfeiffer and Pfeil, 2018).

We note in conclusion that further optimization of dropout techniques may also enrich neuroscience. In particular, the well-described brain regional differences in the density of serotonergic fibers, currently unexplained functionally, might be associated with different levels of plasticity in these brain regions. For example, a high level of plasticity is likely to be beneficial in prefrontal cortical circuits, but such plasticity may be undesirable in brain circuits that control vital organ functions (and may lead to neurological problems). To our knowledge, such analyses have never been carried out. Further insights into dropout algorithms based on FBM, as well as on other anomalous diffusion processes, will strongly motivate this experimental research.

## 6 Author Contributions

SJ proposed that functional similarities may exist between the stochastic organization of brain serotonergic fibers and ANN dropout (as a part of his larger research program) and wrote the first draft of the manuscript. CL implemented the FBM-based dropout, performed the computational analyses, and generated all figure panels (with the exception of Figure 1A-C). ZZ supervised the ANN analyses and edited the manuscript. SJ and ZZ are the Principal Investigators of their respective research programs.

## 8 Acknowledgments

We thank Melissa Hingorani and Kasie C. Mays (members of the Janušonis laboratory) for their experimental work which has contributed to the development of the presented approach.

## 9 Data Availability Statement

The Python and PyTorch scripts used in this study are available at https://github.com/deep-deep-learning/fbm-dropout.

